# Ultra-sensitive measurement of brain penetration with microscale probes for brain machine interface considerations

**DOI:** 10.1101/454520

**Authors:** Abdulmalik Obaid, Yu-Wei Wu, Mina Hanna, William Nix, Jun Ding, Nicholas Melosh

## Abstract

Microscale electrodes are rapidly becoming critical tools for neuroscience and brain-machine interfaces (BMIs) for their high spatial and temporal resolution. However, the mechanics of how devices on this scale insert into brain tissue is unknown, making it difficult to balance between larger probes with higher stiffness, or smaller probes with lower damage. Measurements have been experimentally challenging due to the large deformations, rapid events, and small forces involved. Here we modified a nanoindentation force measurement system to provide the first ultra-high resolution force, distance, and temporal recordings of brain penetration as a function of microwire diameter (7.5 µm to 100 µm) and tip geometry (flat, angled, and electrosharpened). Surprisingly, both penetration force and tissue compression scaled linearly with wire diameter, rather than cross-sectional area. Linear brain compression with wire diameter strongly suggest smaller probes will cause less tissue damage upon insertion, though unexpectedly no statistical difference was observed between angled and flat tipped probes. These first of their kind measurements provide a mechanical framework for designing effective microprobe geometries while limiting mechanical damage.

## Introduction

Microelectrodes implanted into the brain are a critical component of new neuroprosthetic applications and brain-machine interfaces (BMIs), including high-density silicon probes (70 x 20 µm)^1^, syringe injectable electronics (~100 µm)^2^, shuttle delivery^3^, and microwire arrays (<20 µm per wire)^4,5^. As these new probes become more widespread and new fabrication techniques become available, understanding the mechanics of brain penetration and insertion of devices in the 10-100 µm size scale is essential for balancing geometric design, materials strength and tissue damage. The act of mechanical insertion into the tissue is critical for success, as large devices can cause traumatic tissue damage and scar formation, while thin devices may buckle under the loads necessary to penetrate the outer pia membrane^4,6,7^. Microwires have long been studied as a model system recording structures deep in the brain^6–10^ with the general expectation is that smaller wire diameters should yield lower damage, yet actual data for insertion mechanics are sparse; current under active investigations are limited to a few different sizes and almost none in the 10-100 µm probe size range. Initial tissue damage studies with lower-resolution instrumentation were inconclusive about size dependence, with no significant differences found between large (cross-sectional dimensions ~ 200 x 60 µm) and small (100-140 x 15 µm) devices^11^. More recent studies on neural survival discovered significant increases for electrodes with dimensions less than 100 µm^12^, substantiated by work with 8.6 µm diameter carbon fiber electrodes^7,13^.

Here, we performed systematic measurements of brain insertion forces and tissue compression as a function of microwire diameter and tip geometry to guide probe design and minimize damage. Mechanical penetration measurements of these soft, ultra-compliant materials is highly challenging due to the large insertion displacements (mm to cm) before penetration, together with rapid and/or low-force events present during insertion. Compromises typically have to be made either in force sensitivity or temporal resolution; many of the features observed here may have been missed in previous studies simply due to instrumental limitations^6,14–16^. Other studies avoided some of these issues by removing the pia, a thin membrane that acts as the main structural barrier to electrode penetration^17^. Coarse relationships were found between wire size, insertion velocity and insertion force in cortical tissue, yet study of only two wire sizes made it difficult to extrapolate overall trends. In addition, typical surgical procedures do not resect the pia, limiting the benefit of measurements without it^18,19^.

To address these measurement issues we developed a high-performance mechanical measurement system using a modified nanoindentation head as a force transducer. Nanoindentation force transducers are among the fastest, most sensitive force-displacement systems available. However, these instruments are designed with maximum displacements on the order of hundreds of µm^20^, compared to the millimeters of compression required before tissue penetration. This large displacement range was achieved by integrating an entire iNano nanoindentor measurement head onto a long-travel linear actuator, and measuring force and time with the nanoindentor in a zero-displacement mode while the actuator traveled at a constant velocity. The combination of mm displacement range, 3 nN force and 1 ms temporal sensitivity is the most sensitive instrument applied to penetration in the brain and was able to observe important aspects (e.g. pia penetration, microvasculature) of the brain penetration process.

Significantly, we discovered the relationship between probe diameter and both the critical force to initiate penetration and tissue compression was linear with probe diameter, not with cross-sectional area, as might initially be expected. The amount of tissue compression scaled as ~ 4 times the microwire diameter, thus smaller probes required less force to insert and caused less compression of the brain. Interestingly, no statistical difference was observed between flat and angle-polished wire tips, while no compressive penetration event was observed for electrosharpened tips for all wire diameters. The required penetration forces are then compared to the buckling force for different wire diameters and material stiffnesses to provide a guideline for mechanical design.

We chose to examine both a common brain mimic, agar hydrogel, as well as freshly excised murine brains. Agar mimics were developed to have similar compression mechanics as brain tissue^21^, yet since rupture of the surface and continuous material penetration are mechanically distinct from compression, it is unclear if agar is an appropriate model. Agar hydrogels were found to be poor mimics of brain penetration; while overall force magnitudes were similar, the insertion mechanism appears to consist of a sequence of large and small stick-slip events, quite disparate from brain tissue mechanics.

## Results and Discussion

### Penetration force measurement

We developed a high-performance mechanical measurement system to monitor the forces and displacements during penetration of brain tissue and mimics. This apparatus, shown in Fig 1, used NanoMechanic’s iNano InForce 50 indentation head as the force transducer. This system consists of a sensitive 3-plate capacitor transduction system, which can measure forces with nN resolution. Typically the system is operated by measuring voltage and capacitance between the center and two outer plates. As the center plate moves closer to one of the outer plates, the voltage and capacitance increase, translating to a force and displacement. Three-plate capacitor transduction systems can measure displacement with sub-nm resolution but are limited to a working distance of ~ 50 µm, the distance between the outer plates. Instead, we employed a fixed-displacement control protocol, wherein active voltage feedback is used to maintain the center plate in the 3-plate capacitor at *z = 0*. When the indenter head encounters a force, the voltage required to keep the center plate fixed is applied via a closed-feedback loop (1 kHz control rate), providing force measurement as a function of time. The transducer position relative to the tissue is then controlled via a low-noise linear actuator moving 20 µm/s to a depth of 2.5mm. Tungsten microwire diameters of 7.5, 15, 25, 35, 50, 80, and 100 µm were measured, both in Agar and brain tissue.

**Figure 1.**
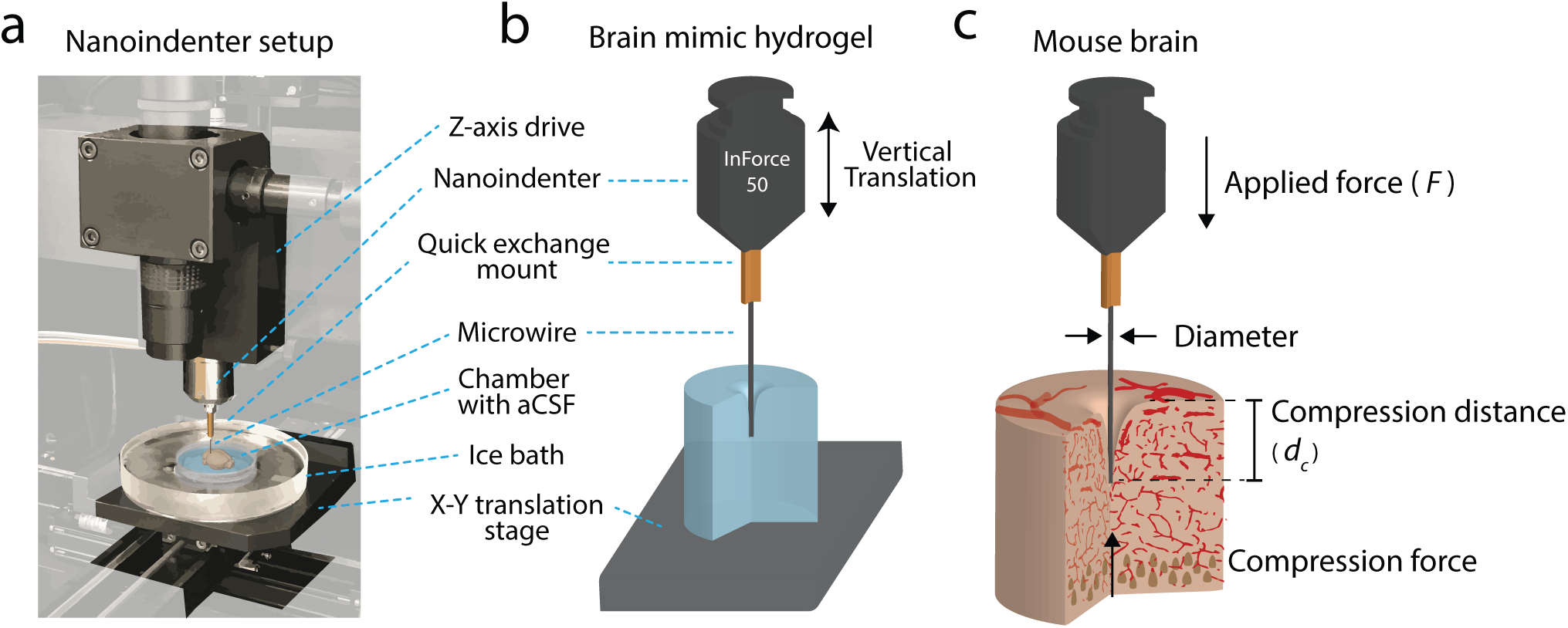
**Ultra-sensitive penetration force measurement apparatus** (**a**) Image of the adapted nanoindenter setup. Background electronics are grayed out to highlight areas of interest. The nanoindenter is mounted on a z-axis drive that provides the vertical translation, and the microwire is mounted onto the nanoindenter via a custom quick exchange mount. Excised brain tissue is placed in a temperature controlled ice bath onto an x-y translation stage. (**b,c**) Schematic illustration of insertion into brain mimic hydrogels, and mouse brain.

### Penetration of agarose mimics

Agarose hydrogels were initially chosen to study penetration mechanics of microwires, due to ubiquitous use as mechanical brain mimics for indentation studies^15,16,21–23^. A representative force-extension curve is shown in Fig 2a for a 40 µm diameter microwire. As the microwire extended at 20 µm/s, the force increased exponentially for the first ~1 mm, roughly ~25 times the diameter of the wire, consistent with previously observed mechanics of highly compliant gels^24^. Visually, as the wire is displaced, the surface of the agar dimples without penetration, until a critical depth is reached at ~ 1 mm. At this critical depth, the wire penetrated through the surface of the hydrogel, and a sudden drop in force is observed. This corresponded with the hydrogel visually relaxing around the wire, with the meniscus approaching its original location, which we interpret as indicating the microwire was inside the gel. From this penetration event both the force at puncture (*F*_*p*_) and the displacement to puncture (*d*_*p*_) were noted (Fig. 2a). A monotonic trend in force as a function of wire diameter was observed, with larger diameter wires requiring higher forces and larger depths to insert (Supplementary Fig. 1).

**Figure 2.**
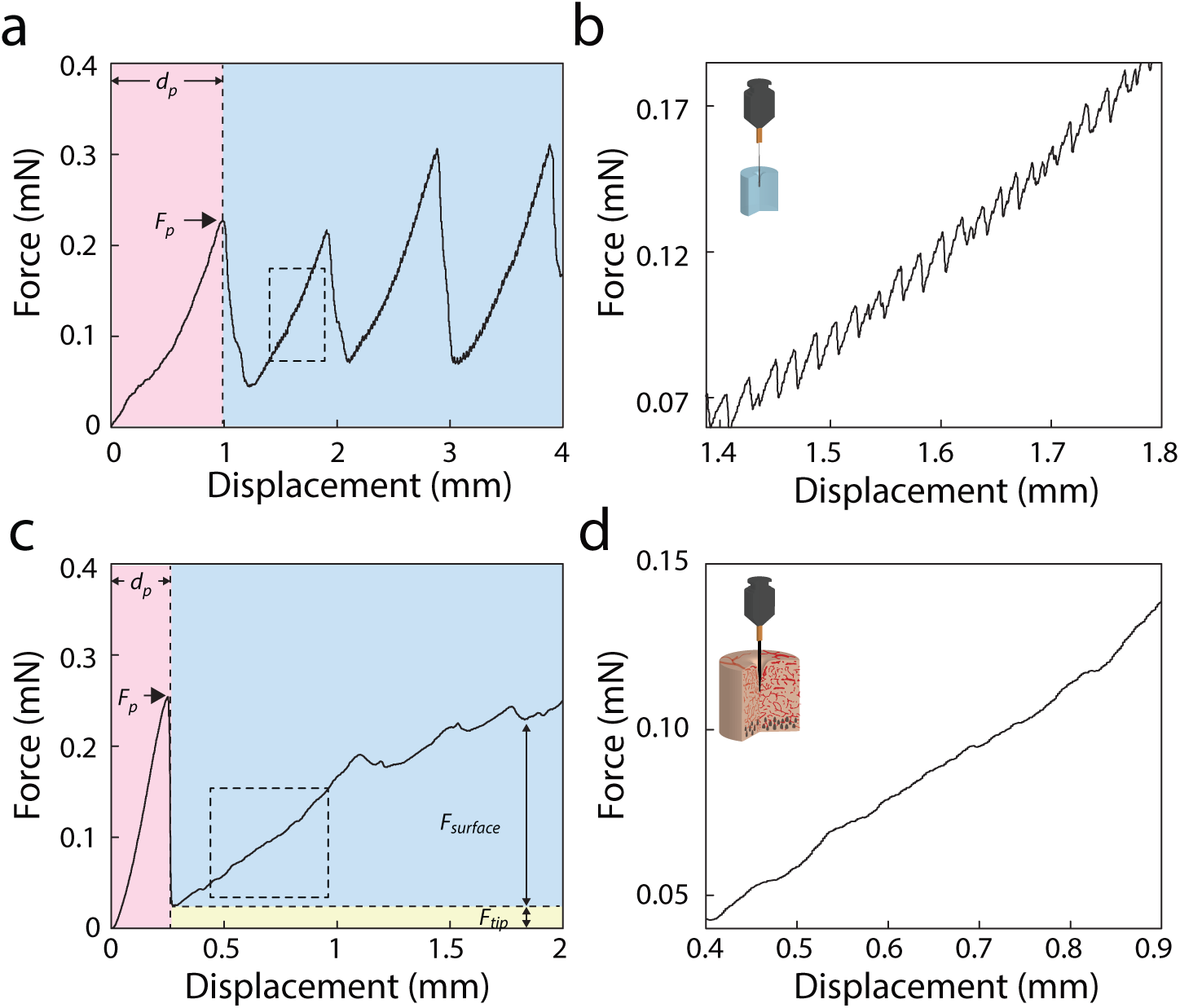
Penetration of brain mimics and excised brain tissue (**a, b**) Force-displacement response for insertion of 40 μm diameter flat-polished wire with an insertion rate of 20 μm/s into 0.6% agarose brain mimic. Penetration into brain mimic and brain tissue is characterized by the sudden drop in force, wherein the puncture force (*F*_*p*_) and displacement (*d*_*p*_) are extracted. Large (mm scale; **a**) and small (µm scale, **b**) sawtooth features in the force-displacement are observed post-penetration, uncharacteristic of previous measurements into brain tissue. (**c, d**) Force-displacement curve for insertion of 15 μm diameter flat-polished wire into acutely excised brain tissue. The force increases exponentially pre-penetration, then increases linearly as a function of depth post-penetration. The linear increase in force is attributed to surfaces forces along the length of the wire (*Fsurface*) in addition to the constant force at the tip (*F*_*tip*_). Excised brain tissue shows different behavior than agarose brain mimics past penetration (**b,d**).

After penetration, distinct saw-tooth features in the force-displacement curves were consistently observed at both µm and mm scales. Large (mm-scale) saw-tooth features displayed the same phenotype as the original breakthrough, distorting the surface of the hydrogel until a dramatic release event occurred, lowering the force and allowing the surface to return near its initial height. We hypothesize these are large scale stick-slip events, as there were no heterogeneities in the material that should lead to such features, and they are highly periodic.

Additionally, similar periodic saw-tooth patterns were observed at the micro-scale (roughly 5 µN magnitude every 15 µm for 40 µm diameter wires), occurring on top of the larger force features (Fig. 2b). Note these did not occur before the penetration event; none were present during the first loading phase. The stick-slip behavior depended on wire size, with larger magnitudes observed with larger diameters (Supplementary Fig. 1b). The characteristic dependence with size suggests a friction-driven series of periodic stick-slip events. No such behaviors have been reported in previous brain mimic or agarose insertion measurements^14,15,25–29^, potentially due to insufficient force and temporal resolution in commercial load cells. Micro stick-slip events could provide valuable information for hydrogel penetration, and demonstrates that high force resolution is necessary to elucidate the underlying mechanisms of insertion into compliant materials.

### Penetration of ex-vivo brain tissue

We then applied the same approach to study the penetration mechanics in freshly excised brain tissue. Using a temperature-controlled bath, the mechanical properties of excised brain has been found to remain similar to live tissue within one hour post-mortem, making it a suitable alternative to testing on live mice^30,31^. Note the dura was removed for these experiments, but the pia membrane was left fully intact. Fig 2c shows a representative force-displacement curve for a 15 µm diameter microwire inserting into the *ex vivo* tissue. Two distinct regimes are observed: tissue compression pre-penetration followed by an abrupt rupture event, and gliding insertion through the tissue post-penetration. The initial insertion behavior consisted of an exponential increase in force as the microwire indented and dimpled the brain surface. At the puncture point near 0.25 mm (*d*_*p*_), the force dropped dramatically and the tissue surface relaxed to near its initial position. The decrease in force was much steeper than observed for any of the hydrogel measurements, consistent with a sudden penetration event.

After entry into the tissue, the force increased approximately linearly as the wire inserted deeper into the tissue (Fig. 2c and 2d). After penetration, the forces acting on the probe are the force of the probe tip displacing the brain tissue (*F*_*tip*_) in addition to the surface forces between brain tissue and the wire (*Fsurface*). As the wire extends, more of the wire surface is in contact with the tissue such that the contribution of surface forces increases. The linear increase in force post-penetration is associated with surface forces along the length of the wire, which we believe to be the dominant force post-penetration. Most traces also had aperiodic variations in the force within this regime, which may arise from heterogeneities in the tissue itself, such as vasculature network or myelinated axon bundles. No large-scale slip-stick events were observed, suggesting the wires glide though the tissue after initial penetration.

The data show that penetration mechanics of brain tissue are mechanistically distinct from agarose mimics, even for materials with similar compression moduli or penetration force. The periodic saw-tooth stick-slip events present in agarose were never observed in any of the *ex vivo* measurements, and the post-penetration force-displacement features were dramatically different between the two. The pia penetration event is also much sharper and abrupt than failure of the hydrogel surface. This suggests that while agarose is a good model for surface compression, it is not suitable to represent penetration and insertion into the material itself.

### Influence of Microwire Diameter

To understand the role of electrode size on the brain insertion mechanics, the force-displacement to insert wires with diameters between 7.5 µm to 100 µm was measured. This size range was chosen due to the increased use of microwires (~ 7 - 20 µm diameter), high-density silicon probes (70 x 20 µm), and syringe injectable electronics (~ 100 µm) of similar scale. Figure 3a shows representative force versus displacement plots of several diameters of flat-polished wires. In all cases the force exponentially increased with insertion depth up to the abrupt pia rupture, followed by a dramatic decrease in loading force, then a linear increase with depth. The slope of the initial loading, indicating the compliance of the brain tissue, was only weakly dependent on wire diameter, with slightly lower compliances for smaller probes (Supplementary Fig. 2). This suggests that over the range of the wire diameters studied, bulk brain tissue mechanics are generally homogeneous, consistent with previous studies on indentation into brain tissue^20,32^.

**Figure 3.**
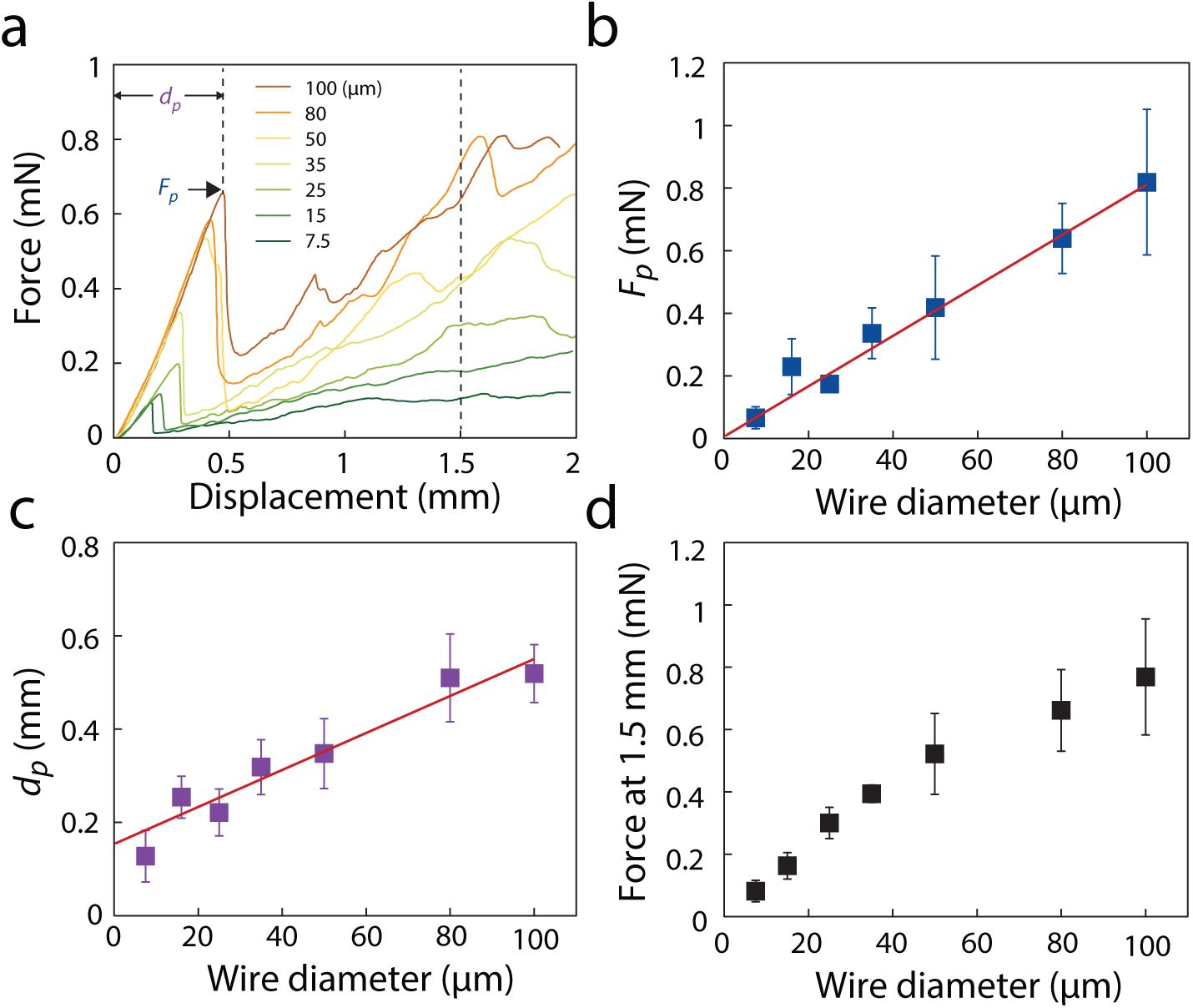
**Influence of Microwire Diameter** (**a**) Representative force-displacement curves comparing the response of different diameter flat-polished probes inserted into the brain at a rate of 20 μm/s. Pia puncture occurs during abrupt drop in measured force, wherein the puncture force (*F*_*p*_) and displacement to puncture (*d*_*p*_) are extracted. (**b**) Relationship between puncture force and microwire is found to be linear. (**c**) The displacement to puncture as a function of wire diameter is also linear. (**d**) The force at 1.5 mm as a function of wire diameter for flat-polished wires.

The pia penetration event shape was highly stereotyped across all sizes, yet the force magnitudes to cause puncture (*F*_*p*_) varied by over an order of magnitude; from 0.066 ± 0.035 mN for 7.5 µm wires, to 0.819 ± 0.233 mN for 100 µm diameter. The *F*_*p*_ scaled linearly with probe diameter, *d*, rather than the cross-sectional area of the tip (Fig. 3b). This observation is consistent with models of crack-initiated or energy-limited failure in compliant materials^33,34^, indicating the failure of the pia likely occurs via one of these modes. From this data, the force needed to puncture the brain could be directly calculated based on the size of the wire:

> *F*_*p*_ *(mN) = 0.00784+(.00807)*d (µm).*

Similarly, displacement to puncture (*d*_*p*_), the distance brain compresses under the device until pia rupture, was also linear with wire diameter (Fig. 3a and 3c). Previous studies have shown that the compression of brain tissue is correlated with brain trauma, suggesting that compression to puncture is an important metric of damage^35,36^. The average compression to puncture *d*_*p*_ had both a constant ~125 um displacement for all sizes, plus a diameter-dependent term:

> *d*_*p*_ *(mm) = 0.127+(4.24)*d*

Thus, while a 7.5 µm wire compressed the brain by ~ 0.1 mm, with a 100 µm wire (similar in size to many Michigan-style probes) compressions of > 0.5 mm are observed, close to 10 -13% of the entire depth of the mouse brain. These results suggest smaller wires are beneficial not just for lower insertion forces, but also lower potential trauma to the tissue prior to penetration.

Post-penetration, the force to push the wire deeper into the tissue increased linearly with depth and was highly dependent on wire diameter. The force after pia penetration in Fig 3a increased significantly with wire diameter, as plotted in Fig 3d. From an ideal mechanical point of view, *F*_*tip*_ is constant with respect to insertion depth and the surface force along the length of the wire is expected to be proportional to the circumference, which has a linear scaling with diameter (π*d). This hypothesis is consistent with the data presented in the next section, where we find an equivalent linear force response after penetration for all tip geometries.

### Tip geometry dependence

Changing the geometry at the tip has been a commonly used strategy for reducing tissue compression for penetrating microelectrodes^17,37,38^, which may result in less tissue damage^39^. For example, a sharper edge has been found to reduce tissue compression for Michigan-style probe arrays, effectively reducing the diameter at the tip and significantly reducing tissue compression and puncture force during insertion^40,41^. To elucidate the effects of probe shape on the penetration mechanics of brain tissue at the micrometer scale, the penetration of angle-polished (24º) and electrosharpened probes were studied in comparison to the flat-polished tips (Fig 4a,b). Two representative force-displacement curves for the insertion of 15 µm and 80 µm wires of the three different tip geometries are shown in Fig. 4c and 4d, with scanning electron microscopy (SEM) images of the different tips (Fig. 4b). The force-displacement curves for the flat and angle polished electrodes are very similar, both in shape and magnitude. The force-displacement behavior during indentation (pre-penetration) was similar between different sizes tested (Supplementary Fig. 2b). In addition, flat and angle-polished wires displayed similar behavior with *F*_*p*_ and *d*_*p*_ as a function of wire diameter (Fig. 4e and Supplementary Fig. 3). While this result is surprising, similar results have been reported when comparing the amount of dimpling caused by sharp vs. blunt micromachined silicon probes^38^.

**Figure 4.**
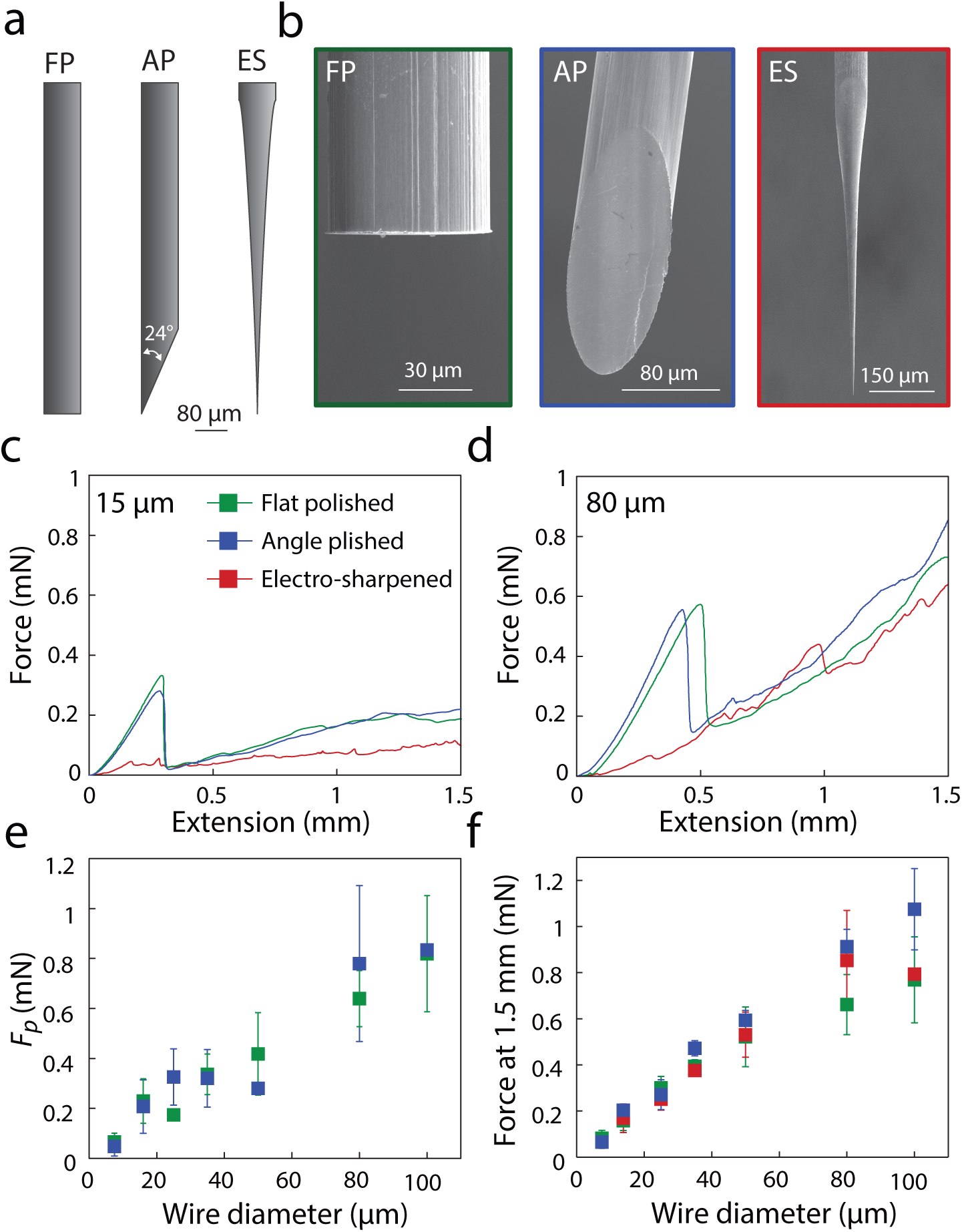
**Tip geometry dependence** (**a**) Graphical illustration of the three different tip geometries studied; flat-polished (FP), angle-polished (AP), and electrosharpened (ES). (**b**) Representative SEM images of the three different tip geometries. (**c,d**) Force-displacement curves for 3 tip geometries: FP (green), AP (blue), and ES (red), for 15 µm diameter (**c**) and 80 µm diameter (**d**) microwires. (**e**) Measured puncture force as a function of wire diameter for FP and AP wires. (**f**) Force at 1.5 mm depth as a function of wire diameter for all tip geometries tested.

In contrast, electrosharpened wires displayed strikingly different behavior, with no discernable penetration event (Fig. 4c,d). There are numerous small humps in the force readout during insertion, but none could be classified as a distinct penetration of the brain surface. Hence, a puncture force was not reported for electrosharpened wires. The compression of the tissue surrounding the wire was also significantly smaller, and no relaxation of tissue post-penetration was observed as with the flat and angle-polished wire penetrations. The forces during initial insertion into brain tissue are dramatically smaller, ranging between 10 - 100 µN, roughly an order of magnitude smaller in comparison to flat and angled tips. Empirically, an electrosharpened tip that has been dulled or bent will still result in a distinguishable puncture event, albeit of much lower peak force magnitude than flat or angle polished microwires. It is possible that the static offset in the relationship between displacement to puncture and diameter diverges in the < 7 µm regime as the size of the wire approaches the size scale of pores in the brain extracellular space (ECS)^42^. Considering that electrosharpened tips have a radius of ~10-30 nm, it could be that the tips slide through the surface of the brain without an observable force event.

After pia puncture, the amount of force to further insert the wire did not depend significantly on probe tip geometry (Fig 4f). No difference was observed between flat, angled, or electrosharpened. This is consistent with the hypothesis that after penetration, the force is dominated by surface forces along the shaft of the electrode and the surrounding brain tissue, rather than crack propagation near the tip.

## Materials considerations for designing brain-machine interfaces

The results from this study suggest that smaller electrodes should reduce physiological damage because of reduced tissue compression. At the same time, research had found that lower material stiffness can significantly reduce traumatic response^7,43^, thus more flexible, soft probes are preferred. Yet as probes become smaller and/or more flexible they may be too compliant to bear the puncture force, *F*_*p*_, and buckle^44^. For example, flexible probes have been inserted through insertion shuttles that provide temporary mechanical stiffness to the probe. Once the probe is successfully in place, the shuttle is removed surgically, or dissolves away^3,4,45^. Unfortunately, these shuttles must be relatively large, which may induce significant primary trauma.

The interplay between the size of the electrodes and a minimum stiffness to pierce the brain without buckling is given by the buckling force threshold, which must be larger than the force to penetrate the brain. *F*_*p*_ was found to be primarily dependent on the diameter of the electrode and is independent of the material modulus. Therefore, the results from this study can predict whether a probe of a given material and size will successfully penetrate the brain before buckling. The theoretical buckling force is calculated using Euler’s formula, given by:

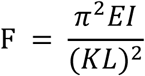

where *E* is the elastic modulus, *I* is area moment of inertia, *L* is the effective length of the wire and *K* is the effective length factor of the wire (2.0 when one end is fixed and the other end free to move laterally).

Figure 5a plots the buckling force as a function of probe diameter for a variety of materials commonly used as brain penetrating electrodes, assuming a cylindrical wire shape of length *l = 3 mm*. This length was chosen because it is deep enough to reach most brain areas in mice, but may need to be larger to be relevant for larger animal models. Plotted on top of the buckling force curves is plotted the measured *F*_*p*_ with probe diameter (red line). Any probe materials or sizes that lie below this line are too compliant and will buckle before penetration, while anything above can penetrate. The minimum wire diameter required for specific materials can be calculated from where the buckling force line intersects with the force to puncture (Fig 5b). For example, stiff tungsten wires can be as small as 12 µm diameter, while gold has to be >20 µm without additional coatings or braces. In contrast, a compliant material like PaC needs to be >60 µm in diameter to penetrate without buckling. Conversely, these larger electrodes will induce increased initial compression in the brain, thus the benefits of softer, compliant materials must be balanced with larger diameters necessary and increased trauma.

**Figure 5.**
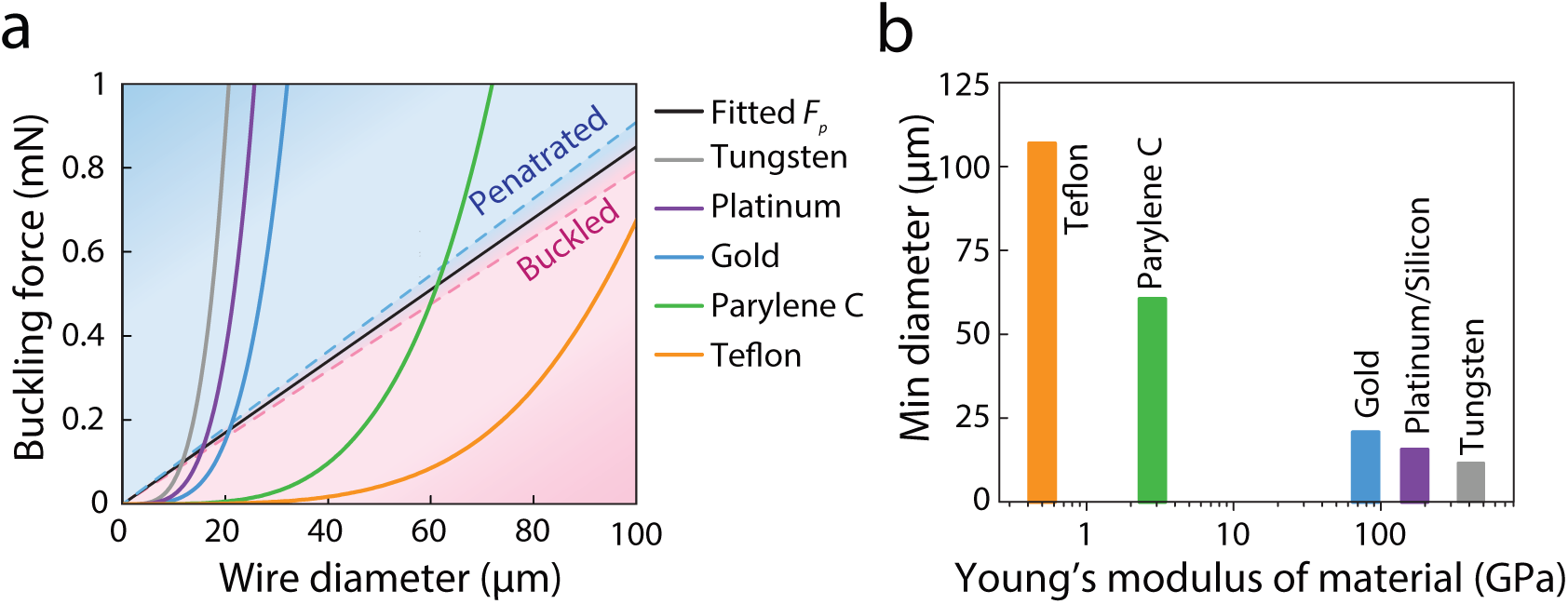
**Materials considerations for designing brain-machine interfaces** **(a**) Buckling force curves as a function of wire diameter, calculated using Euler’s buckling equation. A *K* value of 2 is used, assuming one end is rotationally and translationally fixed (distal) and the brain interfacing end is free. The length is chosen to be *L* = 3.0 mm, a relevant length for brain-penetrating electrodes. Fitted puncture force (*F*_*p*_) based off of flat-polished microwire dataset. Diameters in the blue region predict successful penetration into brain tissue with a wire of a specific material, while diameters in the red region are calculated to buckle before penetration. (**b**) The minimum diameter required for a specific material to penetrate into brain tissue, based off of where the theoretical buckling force intersects with the measured puncture force from (**a**).

## Conclusion

This work addresses the unexplored penetration mechanics of sub-100 µm electrodes by directly quantifying the force and work of insertion into brain tissue with a novel ultra-high resolution force measurement technique. We find significant differences between agar brain mimics and freshly excised tissue. Furthermore, we studied the effects of probe tip geometry on the insertion process and make significant progress towards determining key design criteria for brain-penetrating microelectrodes. Going to smaller probes clearly reduces initial penetration trauma, but more importantly the size dependence is linear, and not quadratic as previous soft solid studies have suggested. This gives some flexibility in terms of geometrical considerations, but nevertheless, there is a 10x difference between 10 and 100 µm probes. These results provide a new basis for determining key design criteria for brain-penetrating microelectrodes.

## Methods

### Instrumentation

We developed a high-performance mechanical measurement system. This apparatus, shown in Fig 1, used NanoMechanic’s iNano InForce 50 indentation head as the force transducer. A displacement control protocol is used to fix the center plate of the iNano, while a low-noise linear actuator moves the indenter head 2.5 mm into the brain at 20 µm/s. The surgical apparatus positions the iNano head above the tissue, then extends the indentation head to push the microwire into the material while measuring force and displacement. A custom program was written to automate force detection of the thin water layer maintained above the hydrogel/brain surface. This was done by vibrating the nanoindenter tip and detecting a phase angle change induced by contact with the water, amplified at the interface by the capillary force pulling onto the tip. This point is referenced to be zero. The system was then programmed to insert by a user specified amount into the hydrogel/brain by a speed capped at 50 µm/s to prevent damage to the tool and assure accurate force measurements (as dictated by the internal hardware speed of the feedback loop).

### Hydrogel Preparation

An agarose 0.6 % hydrogel was made and poured into glass vials. The concentration of the hydrogel was chosen based on literature findings to best match the elastic modulus of the brain^21,46^, done previously via nanoindentation.^20^ Microwires inserted into the hydrogel were not easily cleaned, and typically disposed of consequently. Attempts were made by placing in boiling water, but subsequent insertions in fresh agarose solutions did not show repeating behavior. Using freshly etched microwires, the behavior was very consistent. During insertion experiments, water was added above the hydrogel solution to ensure hydration during the length of the test.

### Fabrication

Tungsten wires of varying diameters (7.5, 15, 25, 35, 50, 80, 100 µm) were spooled, coated with Parylene C (PaC), and subsequently cut into 1” segments. Briefly, aggregates of microwires were placed into glass tubes, infiltrated with Apeizon black wax W, and subsequently polished to accomplish the desired tip angles (flat and 24º) then then released. Stainless steel rods with a 150 µm inter-diameter (ID) bore were cut via electron discharge machining (EDM) to ensure no burr existed after the cut. The parylene coating also acted to increase microwire diameter, allowing for each microwire to be coated with the needed amount to have a final OD of 140 µm, greatly reducing the deviation of each microwire as it is inserted into the 150 µm diameter bore of the stainless-steel rods. Once inserted into the rods, each wire was glued in place using EpoTek 301. Etching in oxygen plasma etched the PaC to expose a length of microwire but kept the PaC embedded in the stainless-steel tube. This method allowed for consistent minimization of angular deviation, as the nanoindentation head can only measure force in the z-axis. For electrosharpened wires, bare tungsten wires were individually submerged in 0.9 M KOH. 2V was applied between the wire and a Pt wire counter electrode using a Keithley 2600 SMU and current was recorded. Etching stopped when the submerged part of the wire broke away from the rest of the wire. This break is observed visually and confirmed by a sharp decrease in current between the two electrodes. Sharpened tips were immediately cleaned with deionized water and isopropyl alcohol and stored for safekeeping.

Every microwire produced was imaged via SEM and documented for quality and reproducibility after manufacturing and each insertion. Cleaning of the tips post insertion was done in enzymatic soap, followed by acetone, isopropanol, ethanol, DI water, and PBS.

### Animal and *ex vivo* brain excision

Adult (10 month) C57BL/6J mice (JAX# 000664) were used for this study. All procedures were approved by Stanford University's Administrative Panel on Laboratory Animal Care. Animals were anesthetized with isoflurane and decapitated. The brain was exposed and chilled with ice-cold artificial CSF (aCSF) containing 125 mM NaCl, 2.5 mM KCl, 2 mM CaCl_2_, 1.25 mM NaH_2_PO_4_, 1 mM MgCl_2_, 25 mM NaHCO_3_, and 15 mM D-glucose. Freshly excised mouse brains were maintained in ice-cold aCSF, and all measurement was done within 1 hour after excision. The mechanical properties of excised brain has been found to remain constant within one hour post-mortem as long as temperature is controlled^30,31^. Repeated measurements using a 25 µm diameter wire showed deterioration of force required to insert past the pia beyond 1 hour after excision. As such, use of a quick-exchange system was developed to allow for as many insertions as possible within one hour of beginning the *ex vivo* preparation. Excised brain was stuck on a petri dish with medical grade cyanoacrylate and filled with aCSF cooled externally in a ice bath (Fig. 1). Surface blood vessels were avoided when possible. Prior to each insertion tests, a thin layer of aCSF was applied to the brain surface to prevent drying. Insertions were all into the motor and sensory cortex areas. Between six to eight insertions were done on each brain, and the position of each subsequent insertion was shifted by 500 µm. The nanoindenter was positioned ~200 µm above the surface of the brain and inserted at 20 µm/s to a depth of ~2 - 2.5 mm. The nanoindenter was pulled out rapidly (1 mm/s) to speed up each insertion and consequently how many wires we could test per brain within 1 hour after excision. Wires were cleaned with enzymatic soap and IPA after each insertion to ensure all tissue residue is removed. To ensure reliability of measurements, subsequent brains tests altered order of diameters used. No clear difference in pia puncture behavior was shown between the two sequential orders tested in the time allotted (Supplementary Fig. 4), suggesting the mechanical properties of the brain tissue remained stable through the recording periods. The large spread seen in these measurements is likely due to variations in blood vessel density or laminar structures of the mouse cortex, even though large vasculature was avoided by visual inspection through a stereoscope.

## Supplementary Figures

**Supplementary Figure 1.**
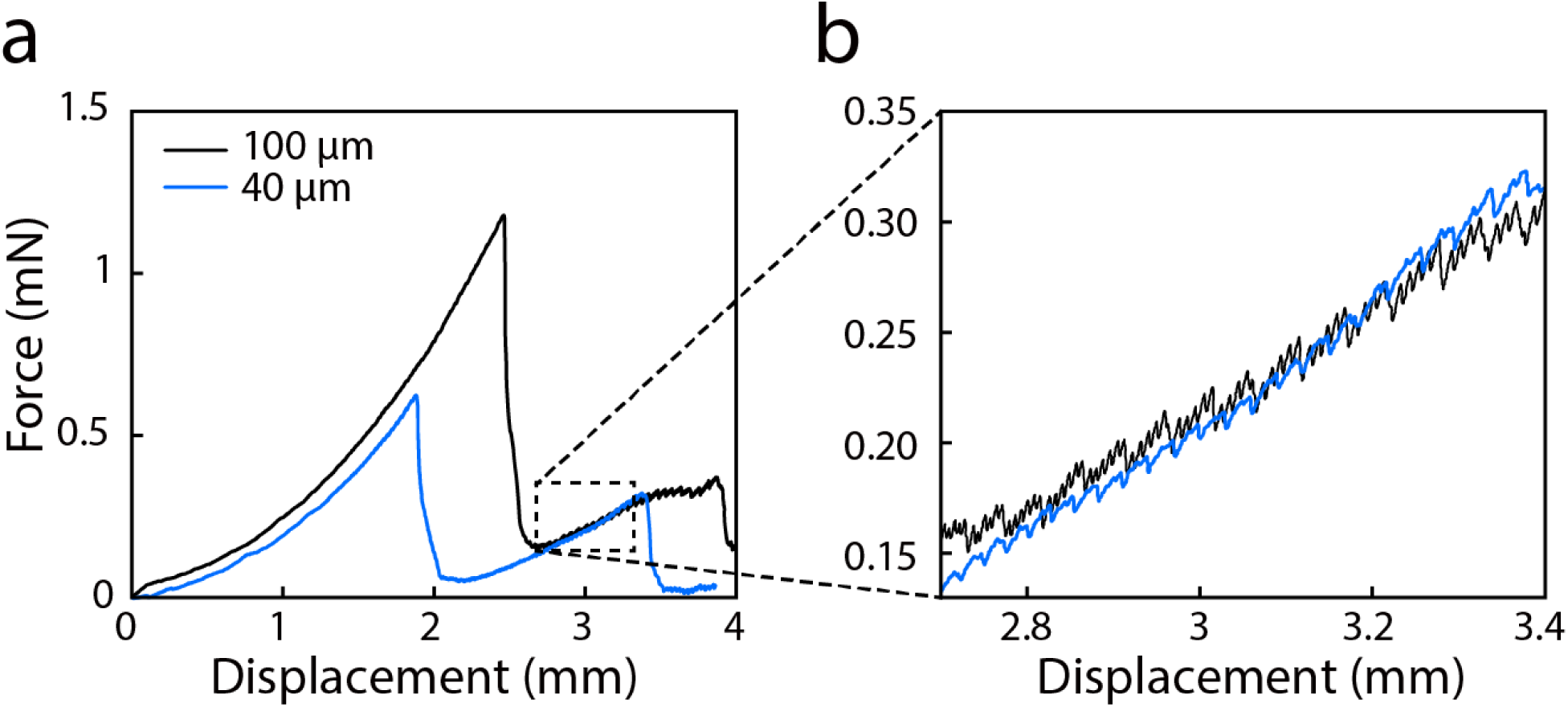
**Size-dependence for agarose brain mimics** (**a**) Representative force-displacement curves for 100 µm and 40 µm flat-polished wires into 0.6% agarose brain mimic hydrogel. (**b**) Zoom-in panel of the post-penetration phase shows differences in the saw-tooth behavior between 100 and 40 µm wires.

**Supplementary Figure 2.**
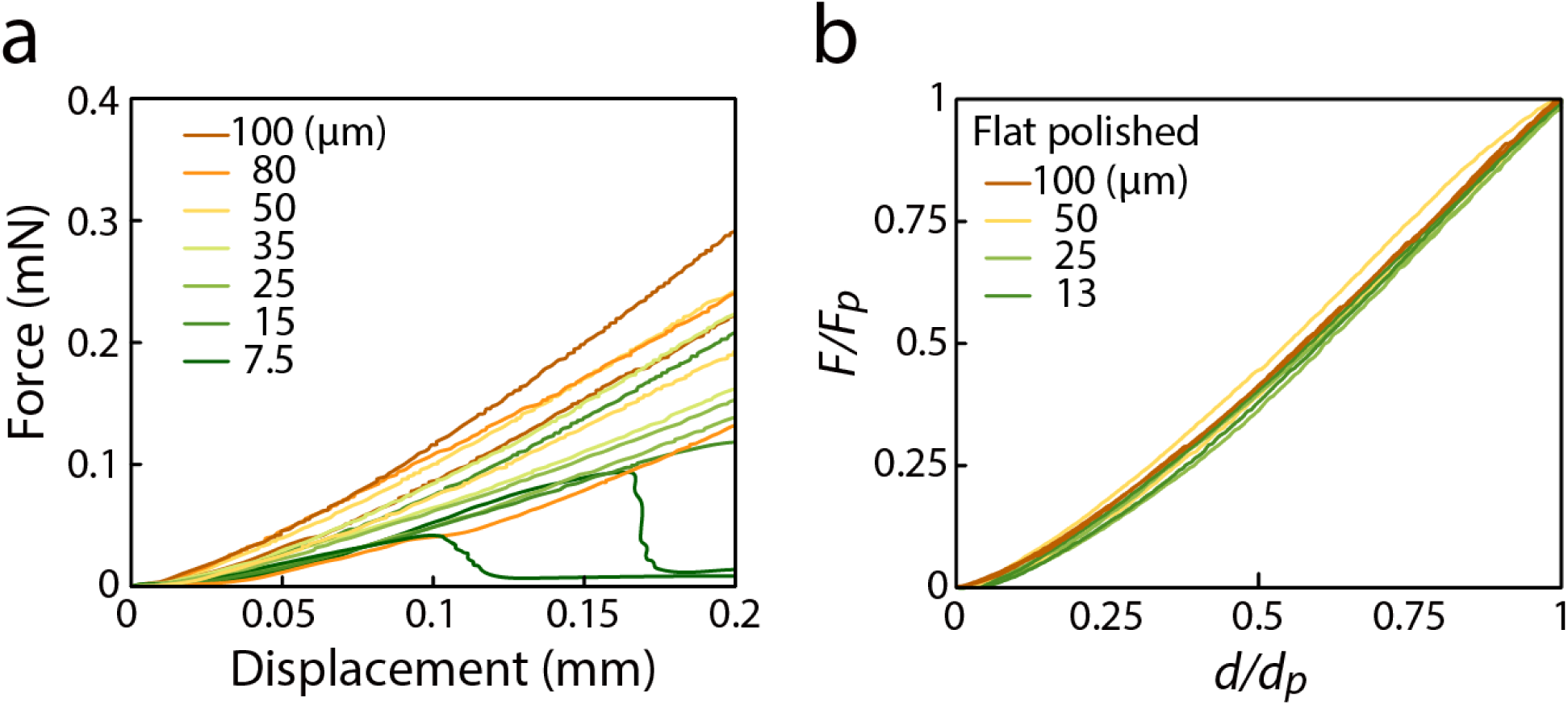
**Pre-penetration curves for flat-polished wires.** (**a**) Two representative raw traces of each size are shown. Coarse differences are found between larger and smaller wires in the force-displacement behavior pre-penetration. (**b**) Pre-penetration curves for flat-polished wires, two of each size, normalized by the force and depth at puncture. No trend with size is observed, suggesting that the underlying mechanics of indenting into brain tissue is similar between 10 – 100 µm.

**Supplementary Figure 3.**
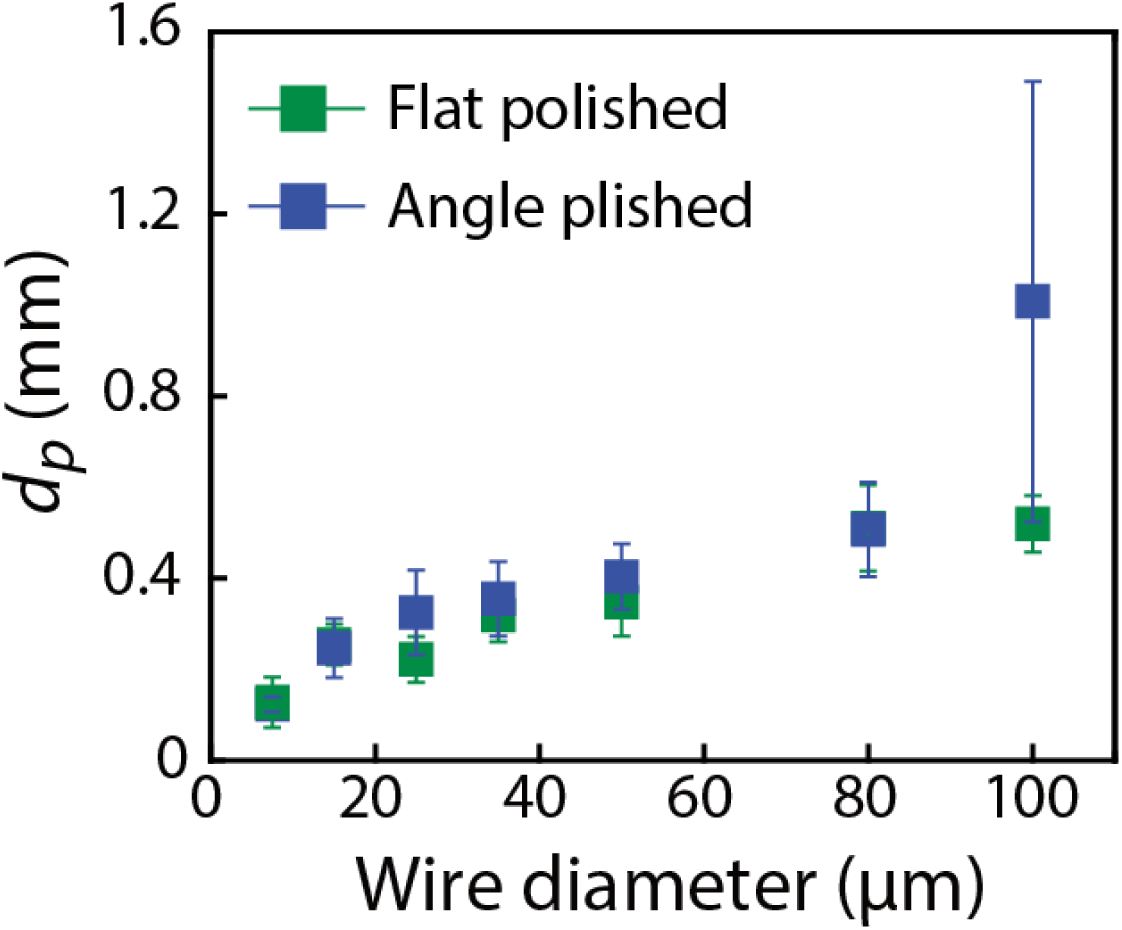
**Tip geometry does not affect displacements to puncture** Average displacements to puncture as a function of wire diameter for flat-polished (green) and angle-polished (blue) wires.

**Supplementary Figure 4.**
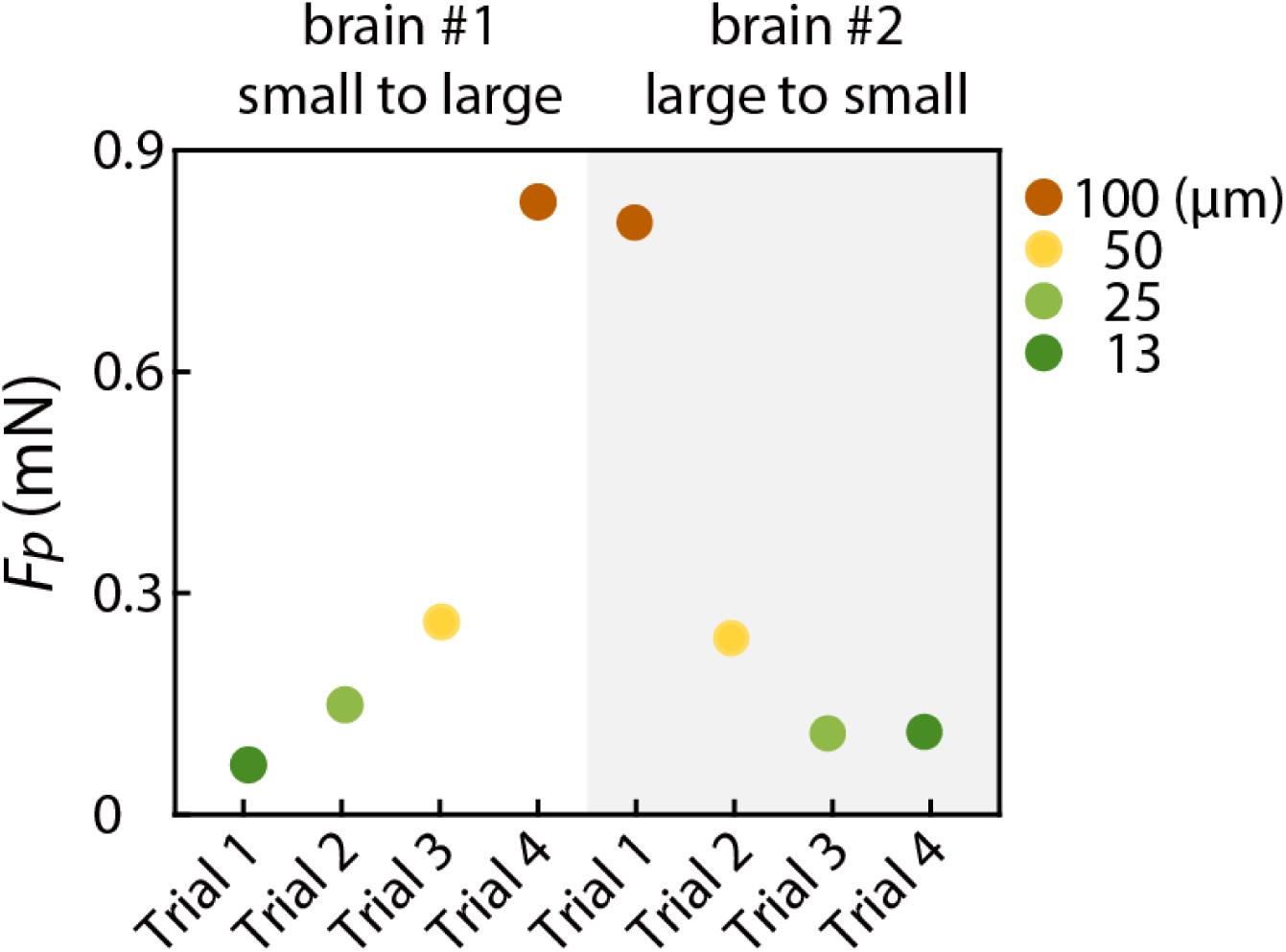
**Reliability of force measurement in *ex vivo* brain tissue** In one experiment, two brains were consecutively tested with reverse order of microwire insertion, from small to large diameters, then large to small. The puncture forces (*F*_*p*_) are shown to be insensitive to the order in which they were tested. These results are consistent with previous work on excised brain in different species^20^.

